# Morphological variability may limit single-cell specificity to electric field stimulation

**DOI:** 10.1101/2023.06.30.547168

**Authors:** Daniel Trotter, Aref Pariz, Axel Hutt, Jérémie Lefebvre

## Abstract

Non-invasive brain stimulation techniques are widely used for manipulating the behaviour of neuronal circuits and the excitability of the neurons therein. While the usage of these techniques is widely studied at the meso- and macroscopic scales, less is known about the specificity of such approaches at the level of individual cells. Here we use models based on the morphologies of real pyramidal and parvalbumin neurons from mouse primary visual cortex created by the Allen Institute for Brain Science to explore the variability and evoked response susceptibility of different morphologies to uniform electric fields. We devised a range of metrics quantifying various aspects of cellular morphology, ranging from whole cell attributes to net compartment length, branching, diameter to orientation. In supporting layer- and cell-type specific responses, none of these physical traits passed statistical significance tests. While electric fields can modulate somatic, dendritic and axonal compartments reliably and subtype-specific responses could be observed, the specificity of such stimuli was blurred by the variability in cellular morphology. These null results suggest that morphology alone may not account for the reported subtype specificity of brain stimulation paradigms, and question the extent to which such techniques may be used to probe and control neural circuitry.

**Author summary:** Over the last several decades there has been increased interest in the efficacy of non-invasive brain stimulation, particularly in determining the limits of specificity of such techniques. Despite this growing area of research, much remains unknown about the interactions of non-invasive techniques with neurons at the single-cell level, notably the importance of morphology to these interactions. We make use of detailed single-neuron models and simulate them in a uniform electric field and demonstrate that the high variability in neuron morphologies may limit how specifically single neurons can be targeted non-invasively. We confirmed this for neuron morphology characteristics at macro- and meso- scales and at varied orientations. Our work suggests that previously reported subtype specificities in non-invasive frameworks are not accounted for by considering only morphological factors.

## Introduction

Non-invasive brain stimulation (NIBS) paradigms are used for a variety of purposes, including attempted treatment of several psychiatric conditions (e.g. major depressive disorder [1, 2], schizophrenia [3], bipolar disorder [4]), and investigating neural oscillation properties and controllability [5–7]. Despite reported clinical efficacy, many questions remain about the mechanisms involved [8] as well as how to optimize their use. The high variability in brain response to NIBS protocols represents a major limitation in these efforts. Such variability has been attributed to genetic and neurophysiological factors, but evidence suggests brain state during stimulation may further impact the response [9–11]. Many contributing factors to response variability are immutable (e.g. subject age, skull thickness, etc), but others, such as the placement of the apparatus (e.g. electrodes or coil), are malleable [10, 12]. The immense combination of free parameters in these protocols greatly complicates parsing which parameters are crucial. This has led to the development of a variety of computational models in efforts to improve the efficacy of experimental approaches and our overall understanding of NIBS [7, 13].

Despite widespread utility, comparatively little is known about the effects of NIBS at the level of single neurons [8]. It is known experimentally, although not widely investigated, that NIBS techniques can modulate the activity of individual cells [12, 14, 15]. Along with this awareness has come an acute interest in parsing the limits of *specificity* of these techniques. That is, the extent to which neuron traits (e.g. subtype, morphology) govern the ability of a single NIBS protocol to acutely activate or suppress the response of neurons sharing similar characteristics. Preliminary efforts have been made to elucidate a relationship between response variability and neuron morphology via intermediate morphological models (that is, models with a limited set of stereotyped, structural components) with nominal success. One such simplified model examined the relationship between many physical traits (e.g., branching, compartment widths and lengths, etc.) of a basic neuron and found, on a trait-by-trait basis, they alter the strength of electric field (E-field) necessary for the neuron to fire [16]. Moreover, even for the most simplified case of a straight-cable neuron model [17], changing a single property, such as the length of the cable, was found to influence the response. Collectively this supports the consensus that physical morphology acts as an avenue for specificity within NIBS frameworks.

Experimental work in alert non-human primates, at field strengths typically applied to humans, has shown transcranial electrical stimulation to reliably control the timing, but not firing rate, of spikes in individual neurons [14]. However, that experimental work found no notable difference in phase locking between pyramidal (PC) and parvalbumin (PV) cells, a result at odds with what has historically been expected of individual neurons, which are long theorized to have subtype- and layer-specific differences in NIBS response [7, 18, 19]. In light of this, it may then be purported that other neuron traits (e.g. morphology) that vary widely across neuron populations may act as the limit to specificity.

Here, we put these results to the test using detailed morphological multi-compartment models from real mouse neurons to computationally study the effects of NIBS on evoked responses of neurons of different types (PV, PC) and populating various cortical layers via the application of a uniform E-field. We specifically examined which, if any, physical characteristics might be involved in generating cell-type and/or layer-specific responses. To do so, we developed metrics quantifying various aspects of cellular morphology, both at the level of whole cells and the compartments they are modelled from, ranging from cell volume to compartment lengths, diameters, number of branches, cellular orientation as well as interneuron myelination [20–22]. Using field strengths commensurate with those used in non-invasive experimental settings, we characterized how evoked responses varied as a function of each of these metrics, linking single-cell morphological characteristics to response specificity. Axonal, dendritic, and somatic compartments could be modulated reliably through changes in E-field amplitude and orientation, and both subtype and layer-specific responses could be observed. Yet, none of these results could be attributed to morphology: the aforementioned metrics didn’t show any statistically significant influence in supporting subtype- and/or layer-specific responses. These results suggest that the interplay between stimulation parameters and physical neuron characteristics is complex in its direction of neuron excitability and that other biophysical features, such as ion channel density and membrane time constant, might be more relevant. The abundant variability of these traits across and between neuron subtypes [23, 24] may limit the specificity of non-invasive stimulation paradigms.

## Materials and methods

### Neuron models

The models used here were created by the Allen Institute for Brain Science based on the morphologies and biophysical properties of real neurons found in the primary visual cortex of mice [20, 21]. That is, these computational mathematical models of single neurons were created based on slice electrophysiology and morphology reconstruction data. A primary objective of the Allen Institute in developing these models was to incorporate active dendritic conductances due to their importance in the input-output relationship of cortical neurons [20]. This was accomplished by including active, Hodgkin-Huxley nonlinear conductances in the dendrites. For the *excitatory* neuron models there are four separate compartment types incorporated: axonal, somatic, basal dendritic, and apical dendritic. In contrast, the *inhibitory* neuron models only use three types of compartments: axonal, somatic, and dendritic. For both model types active and passive properties are considered. For each model, the Allen Institutes packages provide an assembly of MOD files that implement the respective mechanisms for the different compartment types using the biophysical parameters (e.g. conductance, decay time constants, etc.) as measured from the individual mouse neuron. There are 26 free parameters in each model: 18 active conductance densities, 4 intracellular Ca^2+^ dynamics parameters, and 4 passive parameters; specifics of these parameters can be found in the Allen Institute’s databases [20]. Passive parameters are set to a singular value and uniformly distributed across all compartments in a given cell model. For active channel mechanisms, the properties change between the different compartment types (e.g. somatic, axonal, dendritic).

For the purposes of the simulations done in this work, two types of models were considered: those based on pyramidal cell (PC) morphologies and those based on parvalbumin cell (PV) morphologies. A summary of the number of models used from each type and layer is shown in Table 1. All simulations were conducted in a mixed Python - NEURON environment, with the models having been created in a NEURON environment [20, 25, 26]. Subsequent data analysis was performed using the NumPy, Matplotlib, Seaborn, SciPy, and Pandas Python libraries [27–31].

**Table 1.** Summary table of the number of morphological models (No. morphs) for each cell type and cortical layer used in the simulations performed here. Pyramidal cells are PC and parvalbumin cells are PV.

| Cell type | Cortical layer | No. morphs |
| --- | --- | --- |
| PC | 23 | 3 |
|  | 4 | 5 |
|  | 5 | 5 |
| PV | 23 | 5 |
|  | 4 | 4 |
|  | 5 | 5 |

In the NEURON environment, the membrane potentials of the models are calculated using the cable equation:

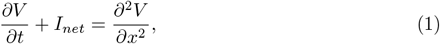

where *I*_*net*_ and *V* are the net current (ionic and injected) and membrane potential, respectively. This is then approximated to its spatially discretized form such that the neuron is reduced to a set of connected compartments, and Eq (1) becomes a family of equations,

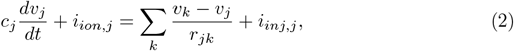

where *c*_*j*_ is the membrane capacitance of compartment *j, i*_*inj,j*_ are the injected currents, and *i*_*ion,j*_ includes all currents through ionic channel conductances. The right-hand side of the equation is the sum of the axial currents that enter the compartment from its adjacent neighbours. A detailed expansion on this derivation can be found in chapters 3 and 4 of the NEURON book [32].

Within this framework, any spatial variation in the membrane current is approximated as its value at the center of a given compartment. Within NEURON’s framework, compartments of the same size are grouped together as a section which contains all of these compartments as segments of the section [25, 32]. This is done for computational efficiency, as Eq (2) may then be re-formulated as,

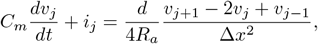

where Δ*x* and *d* are the compartment length and diameter, respectively. If these are the same between compartments, as is the case for a section, then *R*_*a*_Δ*x/π*(*d/*2)^2^ is the axial resistance, and *C*_*m*_*πd*Δ*x* is the compartment capacitance.

The electric field is applied to all compartments in a cell using NEURON’s extracellular function [25]. This function adds the field in millivolts such that it is connected in series with the conductance of the last extracellular layer of the model. For these simulations, we use a uniform electric field in the positive z-direction, such that each compartment receives field *E*_*z*_ = *E* cos*θ* + *ξ*, where *θ* is the angle between the middle of a given compartment and the z-axis, and *ξ* is noise. Ten trials each of subthreshold field strengths *E* = {−50, −30, −10, 0, 10, 30, 50} *mV* were applied, in line with ranges that have been used for similar models [16, 18]. The Gaussian white noise, *ξ*, was added with the field applied to each compartment and is different between compartments and across trials. The cell response was read out from the somatic compartment as a membrane potential and the average response across trials was taken to quantify the specificity of the cell.

### Characterizing neural morphology

In the present framework, the Cartesian coordinates for all compartments in the models are known. This allows for both whole-cell and compartmental characterizations of the morphologies to be made. For macroscopic measures, that is characterizations that encompass the whole cell, we take three quantities 1) the vector magnitude, 2) the length in the *z* -direction, and 3) the elliptical volume. We define the vector magnitude as the mean square root of the sum of the squares for the Cartesian coordinates of all compartments in a given model and calculate it 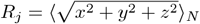, where *N* is the number of compartments in model *j*. For the *z* -direction length (referred to hereafter as length), the value *L* is calculated as simply the difference between the maximum and minimum *z*-direction coordinates. Finally, the elliptical volume is calculated by taking the length, and the equivalent maximum distances in the *x* - and *y* - directions, halving each of them and utilizing the formula for the volume of an ellipsoid, *V* = *π/*6(*x*_*max*_*y*_*max*_*z*_*max*_).

Additionally, at the level of whole cells, we sought to investigate the effects of orientation. To do this, the models were rotated about the *y* -axis using a rotational matrix within the field at *ϕ* = 0^°^, 90^°^, and 180^°^. At each orientation, the models were simulated with the same *E* = {−50, −30, −10, 0, 10, 30, 50} *mV* fields. For these simulations, recordings of the stimulation responses were taken for all compartment types to create a more complete polarization profile. Notably, in these simulations the response is recorded at the center of a given section to approximate the value of all segments within the section.

For the more mesoscopic measures, that is those at the level of the compartments that comprise the models, we consider again three quantities: 1) the average compartmental length, 2) the average compartmental diameter, and 3) the number of branches. For each of these metrics, we further separate into the dendritic and axonal measures. These metrics are defined in the model reconstruction files themselves [21, 25], and were subsequently extracted and averaged across morphologies for the same neuron subtype and/or layer.

### Simulations and analysis

For the simulations conducted here, the cells are artificially positioned such that the somatic section, a singular section for all models, is at the origin and all other compartment coordinates are normalized with respect to this compartment. This is done using the *a priori* known Cartesian coordinates for every segment in the models. Additionally, the orientations were taken to be in standard depiction format, that is with the dendritic arbour in the positive z-direction. To improve the biophysical accuracy of the models, myelin was added to the axonal compartments of the neuron models by randomly selecting a fraction of the axonal segments and setting their ionic conductances (and by extension their ionic currents) to be zero. The myelinated segments in a given simulation were chosen from a Bernoulli trial for each axonal segment with a probability of myelination *p* = 0.7 in the PV neuron models and *p* = 0.15 for PC models. These values of myelination probability were chosen in line with the percentage of the axon length expected to be myelinated from experimental work on these neuron types [22].

### Susceptibility

The curves of membrane potential resulting from the application of various applied field strengths as recorded at the somatic compartment in the upright (or 0^°^) orientation serve as a representation of the susceptibility of a neuron to the applied field. Here, we measure this property using the slope of the membrane potential curves:

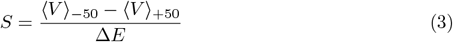

where Δ*E* is the difference between the E-field strengths for the cases (here equal to 100) and ⟨*V* ⟩_*i*_ are the average membrane potentials at field strengths *i* = {−50, +50}. As Δ*E* = −100 here, the resulting values of *S* are also negative.

These susceptibility values offer a metric to quantify the influence of the physical morphology characteristics on the response of the neurons to stimulation by looking at their correlations. Correlations of these susceptibilities are calculated with respect to the different morphological traits using SciPy’s linear regression function, which calculates the Pearson correlation coefficient, *R*, and its corresponding *p*-value using a Wald Test with t-distribution of the test statistic [30]. These correlations then allow us to discern which morphological trait (if any) may contribute to the specificity of the NIBS protocol.

## Results

### Characterizing neuron morphology variability

The neuron models created by the Allen Institute are based on the morphologies of pyramidal (PC) and parvalbumin (PV) neurons from layers (L) 23, 4, and 5 of mouse primary visual cortex [20, 21] (see Methods). Example morphologies of these models from each layer and subtype are shown in Fig 1A. Seeking to investigate the effects of morphology on the response of such neurons to NIBS is a very high dimensional problem due to the high variability seen between neurons. We quantified the variability in physical properties between neuronal subtypes at the level of 1) whole cells (i.e., vector magnitude, length, volume, myelin; see Methods); and 2) compartments (i.e., axonal/dendritic compartment lengths, diameters, branches; see Methods) and evaluated the contribution of each on evoked responses to uniform electric fields.

**Fig 1.**
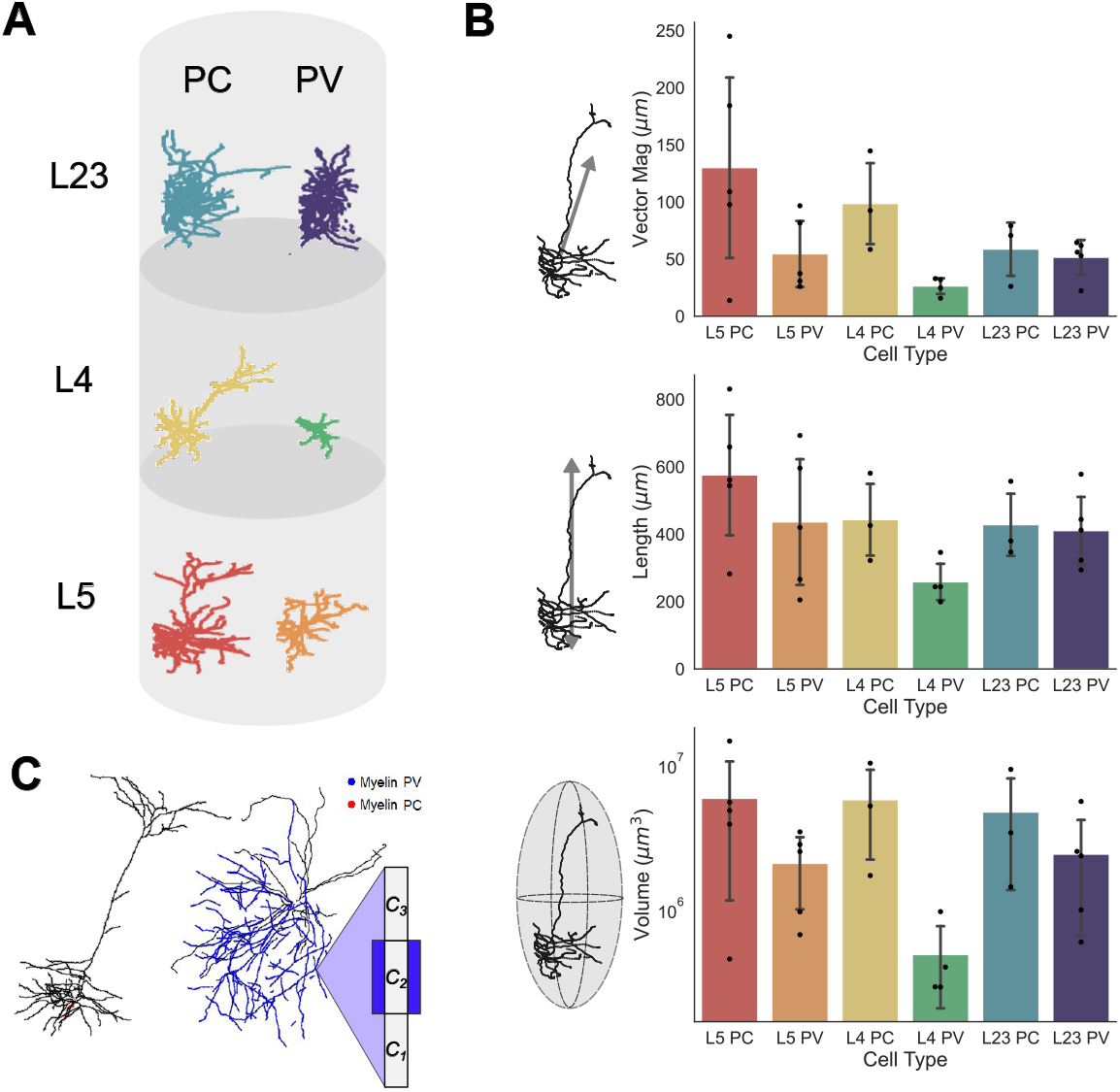
Whole-cell morphological characterization. **A)** Example morphologies of one model neuron each for pyramidal (left) and parvalbumin (right) neurons in layers (L) 23, 4, and 5. **B)** (top) Vector magnitudes of all neuron models split by layer and type; (middle) Length in the z-direction of all neuron models; (bottom) ellipsoid volume occupied by neuron models. Error bars are standard deviations. All bars are colour-matched to the morphologies shown in the example column of panel (A). **C)** Example morphologies and schematic for a PC and a PV neuron model with representative myelination on their axons. In (C), *C*_*i*_, *i* = 1, 2, 3 represent the axonal compartment and the blue sheet over *C*_2_ is myelin.

At the whole-cell level, vector magnitude refers to the mean of the square root of the summed squared cartesian coordinates of each compartment. The length refers to the maximum length of the neuron in the z-direction and volume is the maximal elliptical volume it occupies. Separating the mean vector magnitude for each neuron subtype and layer, the PV neurons generally had smaller vector magnitudes, and so less spread, than their PC counterparts, with L23 having the smallest difference between the two subtypes (see Fig 1B, top). In line with this, the PV neurons were on average shorter than their PC counterparts in the same layers, with L23 having the minimal difference between subtypes (see Fig 1B, top and middle panels). As was the case for the vector magnitudes and lengths of the models, the volumes occupied by neuron subtypes in the same layers were similar with the exception of L4 where the PV neurons occupied substantially less volume (Fig 1B, bottom). Within subtypes, these macroscopic characterizations were reflective of each other, such that longer neurons tended toward a larger vector magnitude and by extension elliptical volume.

Additional to these whole cell characterizations, compartment-level variability may also be considered (see Methods). These can be separated further into their average dendritic and axonal components for each subtype (see Fig 2 top and bottom rows, respectively). This scale of breakdown for the model properties is important as it has been demonstrated that, in isolation, each of these features may affect the field strength required to evoke depolarization [16]. For these models, the compartment lengths show more variability across the layers and subtypes in the axonal compartments than in the dendritic ones (Fig 2A). Further, the dendritic compartments are on average longer than the axonal compartments, however, there are exceptions such as in L5 PC models where axonal and dendritic compartments were around the same length.

**Fig 2.**
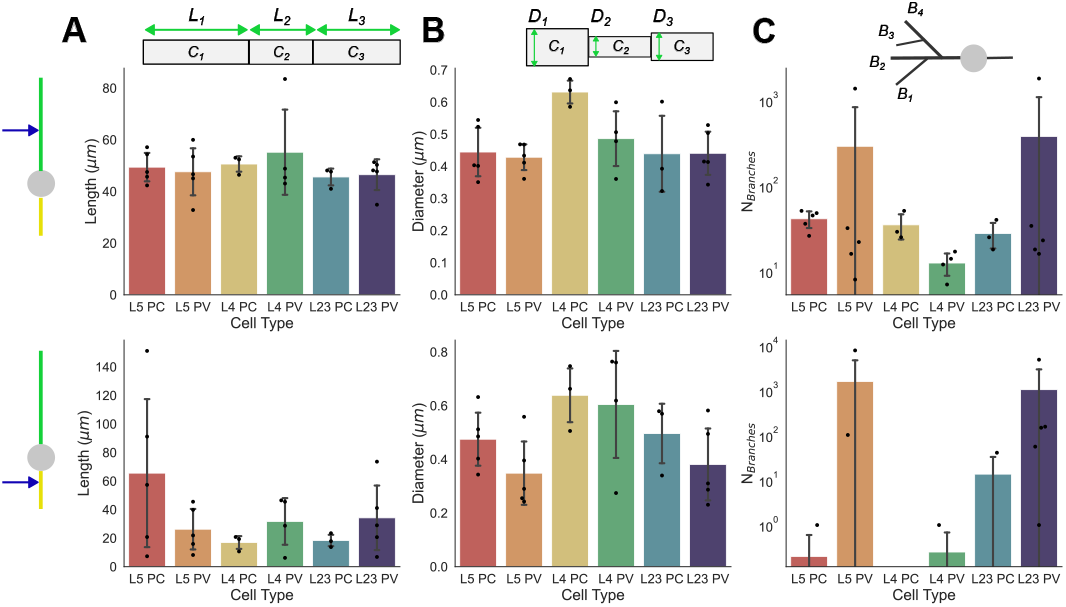
Dendritic and axonal compartment characterizations. Compartmental properties averaged across neuron cell-subtypes of dendrites (top row) and axons (bottom row). Individual points plotted on/above the bars are the compartmental quantity average for individual models; bars are the mean across models. **A)** average compartment lengths (*L*_*i*_), **B)** average compartment diameter (*D*_*i*_). **C)** average number of branches. Error bars are standard deviations.

By contrast, the compartment diameters are relatively similar between the dendritic and axonal compartments (Fig 2B). The diameters observed between subtypes and layers are also, on average, relatively similar to each other. Branching is, generally, observed to be much more prevalent in the dendritic compartments than the axonal ones (Fig 2C). However, there are exceptions to this, such as in the L5 and L23 PV models where the axons are very densely branched. Indeed, the PV models generally have more branching than their PC counterparts in the same layer regardless of whether the branching is considered at an axonal or dendritic level.

### Subtype- and layer-specific evoked responses to uniform electric fields

We first examined the evoked response of these models across layers and subtypes, independently of their morphological variability. To do this, we apply a uniform electric field to the neuron models (Fig 3A). The sign and strength of the field will influence whether the field depolarizes or hyperpolarizes the model at its somatic compartment, hence the relative evoked response may be quantified by averaging membrane potential fluctuations across independent trials (see Fig 3B and Methods section).

**Fig 3.**
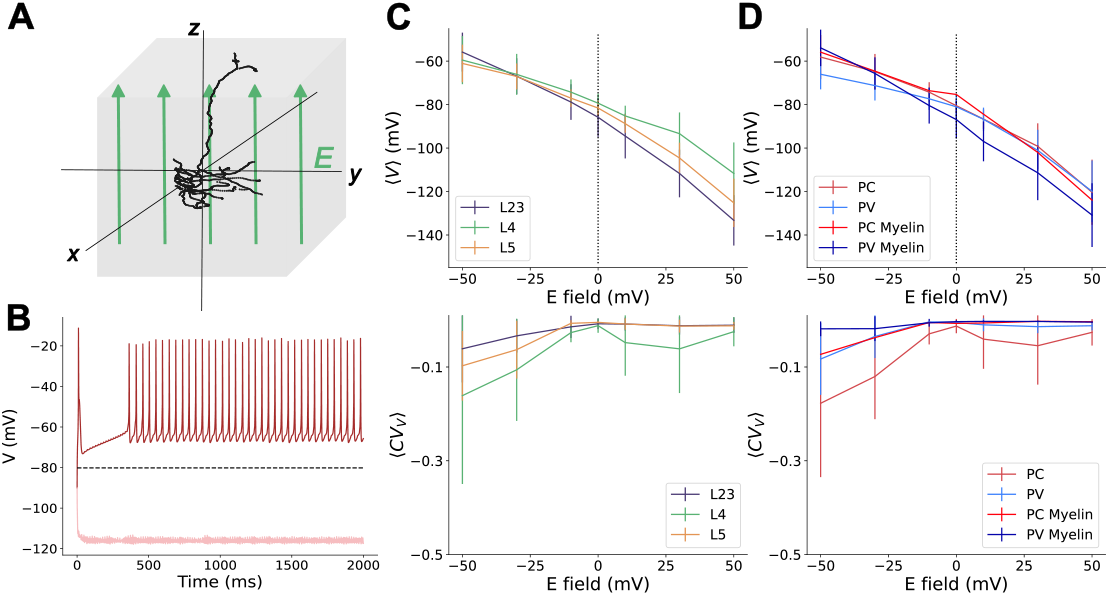
Uniform electric fields impact on somatic membrane potential. **A)** Schematic of uniform electric field, *E*, being applied to neuron model; **B)** Example responses of a PV layer 5 neuron model to a uniform field for *E* = −50mV (dark red), and *E* = +50mV (light red). The black dashed line is the average response to noise (*E* = 0mV) over ten trials. **C)** Top: Average membrane potential across trials pooled by layer. Error bars are the standard deviation. Bottom: average coefficient of variation (CV) of neurons at different E-field strengths pooled by layer. Error bars are the standard deviation. **D)** Top: Average membrane potential across trials pooled by neuron sub-type. Bottom: average CV of neurons at different E-field strengths pooled by neuron sub-type. Error bars are the standard deviation.

While not statistically significant, differences in evoked responses could be observed between the subtypes and layers. Pooling the neurons by layer (neglecting the subtypes) there is no significant difference between the somatic responses observed. The lowest coefficient of variation of these membrane potentials, ⟨*CV*_*V*_⟩, is observed for the L23 models, with the highest being the L4 models (Fig 3C). By comparison, ignoring the layers and pooling the neuron models based on their subtypes, there are visible differences between the depolarization responses of PC and PV cells (Fig 3D). Of note, the differences observed between subtypes manifest differently in the cases where the PV neuron models are myelinated or not. For unmyelinated PV neurons, the same field strength leads to less depolarization than is observed in the PC models at the same field strength. However, in the myelinated PV models, the response to the same field strength eventually catches up with the PC models in terms of depolarization. Interestingly, the myelinated PV models also hyperpolarize more aggressively than either the PC or unmyelinated PV models (Fig 3D). The lack of statistically significant differences observed in these poolings suggests limitations to the extent of specificity attainable via NIBS techniques for circuit control. The average *CV*_*V*_ for the pooled subtypes suggests that the PC models have more variation in their responses. Based on the severity of this curve compared to the PV models and the similarly shaped curves in the same plot pooled by layers, it is likely that the majority of this variation comes from the L4 PC models.

Additional to interrogating the effect of model subtype and layer on their contributions to response variability in NIBS, we also seek to understand whether there are significant impacts from the orientation of the neuron within a uniform field. An example neuron is shown with its polarization profile in Fig 4A, at three major orientations about the y-axis: from left to right, 0^°^ (as was used for the prior analysis), a 90^°^ and 180^°^. From this, we can clearly see that the overall polarization of the neuron is altered by the orientation within the field, however, at the level of the somatic compartment, there is no meaningful change in the depolarization versus hyperpolarization (Fig 4). That is, while rotating the neuron by 180^*o*^, or even 90^*o*^, will result (for this neuron) in a stronger depolarization however, the net response to the applied field is largely conserved at the soma, and do not differ significantly from the somatic readings of the other models (shown in grey). However, it does have a marked effect on the response of a given dendritic or axonal compartment (see Fig 4B middle and right). This suggests that, for neurons in these layers, the attainable specificity through NIBS techniques is insufficient to control neurons with a specific orientation.

**Fig 4.**
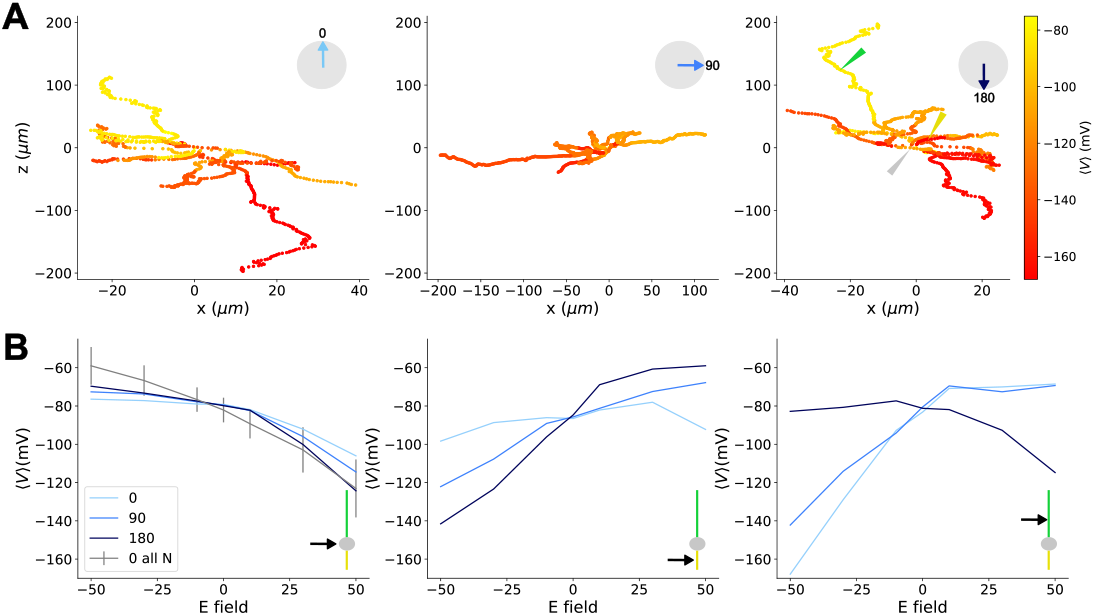
Orientation effects on the polarization profile of neurons. **A)** Average membrane potential per compartment for *E* = −50 applied to an example L5 PV neuron model as it is rotated 0^*o*^, 90^*o*^ and 180^*o*^ (left to right) with respect to the y-axis. Orientation is indicated in upper right of each plot by the arrows on the circles. The lightness of blue on the arrow corresponds to the curves on plots in (B). **B)** The relationship between the example neuron model orientation and average membrane potential with respect to the applied E-field. The simplified neuron schematic in the bottom right of each plot indicates which compartment type is being plotted (left to right: somatic, axonal, dendritic). The exact section of the neuron being recorded from is indicated by the colour-matched triangles in the rightmost panel of (A). In the left panel of (B) the average 0^*o*^ orientation membrane potential from the somatic compartment of all models is indicated by the grey line with the error bars as their standard deviation.

### Examining the relationship between morphological variability and response specificity

Having these response curves as a function of applied field strength, it becomes of interest to ask: what drives subtype- and layer-specificity? Specifically, we sought to determine whether any of the characterized morphological attributes substantially influence the response of the neurons. Returning to the physical metrics used earlier to describe the neuron subtypes and layers (Fig 1B and 2A-C), the relationship of these properties to response can be investigated through their correlation. The trends of individual model responses with respect to field strength here are quantified through their susceptibility, measuring the amplitude of the response to a given change in electric field strength compared to the baseline (see Methods).

As with the characterization of physical traits, this can be broken down into considering both whole-cell and compartmental physical traits. Looking at the slopes agnostically (irrespective of the neuron subtype or layer) with respect to the more ‘macro’ quantities (vector magnitude, length, volume occupied, and the number of branches) all yielded non-significant correlations with the occupied volume showing the highest correlation (Fig 5). At the compartmental level, investigation of the slopes of compartment properties also yielded no significant correlations, with dendritic and axonal branching showing the highest correlation among their respective compartmental attributes. This implies that controlling neural circuits with NIBS techniques is limited in its specificity by the high variability found in morphologies.

**Fig 5.**
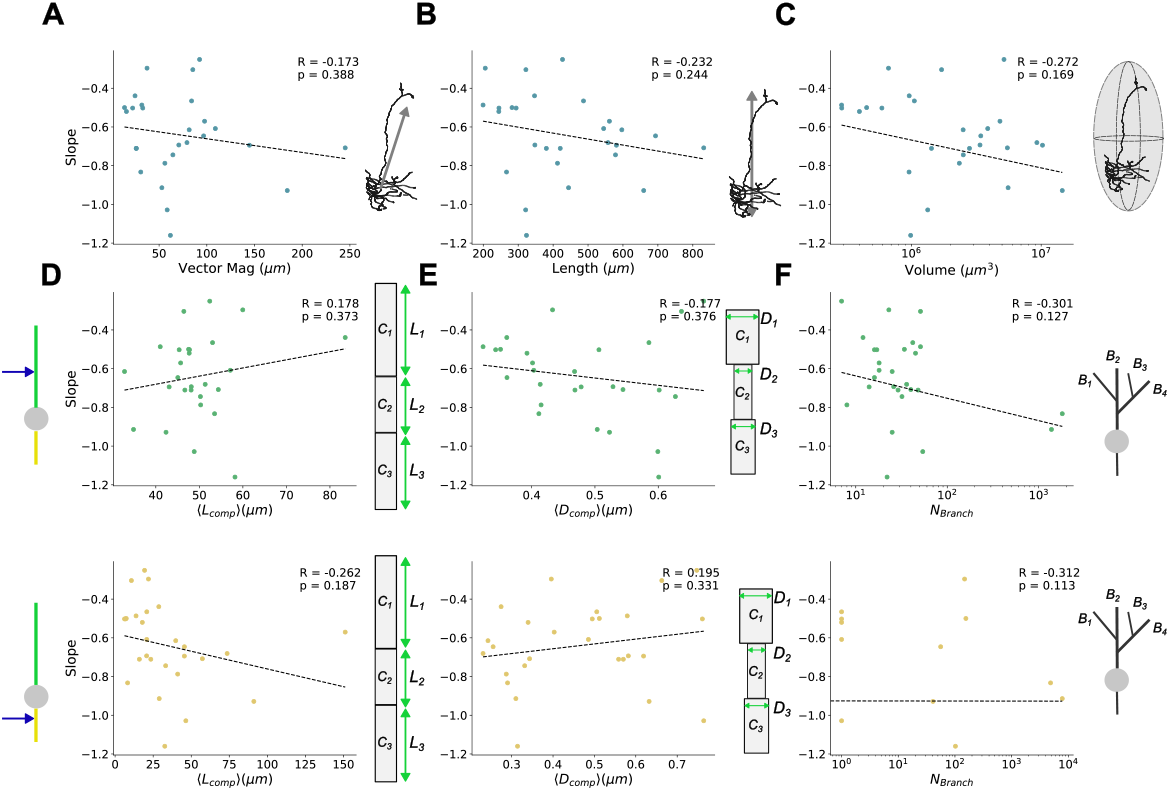
Relating field effects to morphological characterization. Slope of average somatic membrane potential curves (see Fig 3C-D) per neuron as a function of their **A)** vector magnitude, **B)** neuron length, and **C)** volume occupied. At the level of dendritic (upper) and axonal (lower) compartment properties (see Fig 2), the slope of the average somatic membrane potential curves is shown as a function of **D)** average compartment length, **E)** average compartment diameters, and **F)** number of branches. All slopes are calculated from response in the upright (0^°^) orientation. Schematics to the right of each plot correspond to the quantity on their x-axis. The Pearson correlation coefficients, *R*, their significance, *p*, and the line of best fit are found using SciPy’s linear regression package [30].

## Discussion

Previous NIBS studies have demonstrated that single neurons are affected and entrained by low field strengths [14, 17, 33–35]. The extent of specificity NIBS holds over single neurons is less clear and, despite some successes that found no difference in the entrainment of excitatory and inhibitory neurons [14, 15], is largely experimentally intractable. Our computational investigation found minimal difference in the acute specificity of neuron subtypes, and no significant correlation between response variability and physical morphology. These results align with experimental work revealing that highly complex, variable morphologies had no effect on the conservation of physiological waveforms and circuit functions of neurons [36]. Similarly, recent computational work in L5 pyramidal cells has shown morphological variance to be insufficient to reproduce electrical variability [37].

Among the morphological diversities we considered was myelination, which influences neuron response to NIBS [38–40]. The minor differences in the PV, but not PC, models in the myelinated and unmyelinated versions (see Fig 3D) suggest that response variability due to myelination is non-uniform and, consistent with a recent work [33], requires *>*15% coverage to manifest. This may contribute to earlier computational works, without myelination, that found layer-specific differences in the E-field strength required to depolarize neurons in L23 and L56 [18] whereas recent studies involving myelination have shown minimal difference required to evoke firing from L23, L4, and L5 neurons [33, 41]. As the effect of myelination is related to its amount, in brain regions with less myelin, or shorter axons, inhibitory neurons may respond more similarly to their unmyelinated equivalents (see Fig 3D), and hence be less depolarized than excitatory neurons. Indeed, such effects may explain differences in our results and other myelinated models that predicted specific activation of cells across subtypes and layers. Where those studies use morphologies from multiple regions of the brain [19], they may indicate that morphologies across brain regions differ more substantially than those found in the same region, and are hence more specifically selected. Such a distinction may also explain discrepancies between those works and the lack of significant difference found between subtype responses in experimental protocols [14].

In addition to layer- and subtype-based neuron groupings, we sought to explore the influence of physical morphology on neuron response (see Fig 5). Compartment-level studies of simplified neuron models previously found individually, varied physical traits which influence the field strength required to evoke firing [16]. Based on those findings, one could infer that an ideal neuron (that is, one with the lowest required field strength required to fire) would have long, thick dendrites with many branches that disperse minimally from the primary axis the applied field lays on. Moreover, an ideal neuron would have long, narrow axons with minimal branching that disperses widely from the field axis. For the models considered here, there is no class of neurons that fits this ideal (e.g. long but thicker axons; see Fig 2). In line with this, our simulations did not demonstrate any singular physical trait to be the driving source response specificity to the same stimuli between neurons. This suggests the difference between individual neuron models and their response is likely the result of a convolution of the physical properties rather than individual traits determining the outcome.

For computational efficacy, many models of NIBS protocols make use of mean-field type approaches, whether for the whole model population [42] or for subtypes within the population [7]. If groupings of neurons could be derived on the basis of physical morphologies, this would imply the need for adaptations to these models to include more extensive morphological features. In contrast to this, here we have found no significant relationship between susceptibility and morphology (Fig 5); this suggests support for the assumptions made in designing such computational models to not include specific morphology in their frameworks.

## Limitations

Our morphologies come from neurons found in the primary visual cortex of mice; importantly, this limits the extrapolation of the results of these simulations to any expectations for human cells, as it has been shown that a number of neuron properties do not scale from non-human to human neurons [43, 44]. In line with this, recent results showed human L5 PC neurons have unexpected biophysical properties for their size, despite the composition and allometry of the human cortex scaling relative to the nine other species tested [44]. Further, as these models are simulated in isolation, they disregard any potential influence from conductive tissue, distance-to-skull or other cell interactions that may bolster response through biophysical mechanisms, such as ephaptic coupling [13, 34, 35, 45]. White noise is added to account for some of these effects, however, the net efficacy may still be impacted. Additional model-specific limitations can be found in the Allen Institute technical papers and documentation [20, 21].

The neuron models were predominantly simulated in one orientation in free space. However, the overall effects of orientation were considered by rotating the neuron models 0^°^, 90^°^ and 180^°^ about the y-axis and recording from the somatic, axonal and dendritic compartments in all three orientations (see Methods and Fig 4). The observed effect of this change is relatively small at the somatic compartment, although the overall polarization of the model changes. NIBS studies of biophysical models found this minimal effect results from the axon branching and myelination, which drastically reduces the effects of orientation on activation threshold [33]. The layers considered may also reduce the effect of orientation, as L1 and L6 have previously shown more variability in preferential orientation [8, 41]. The average membrane potential in response to orientation changes is more attenuated at the soma than the dendrites and axons, which are two to three times more susceptible to polarization at their terminals than somas [46].

Importantly, for the model neurons, there is no distance component to the applied field as it is applied as directly to the surface of the neuron (see Methods). For real neurons receiving input from outside the skull and encircling brain tissue, the minimum field strength required to facilitate depolarization may be greater than in these models where the field is applied uniformly. Alternatively, having connections to other neurons to bolster the response may also allow *in vivo* neurons that would not individually react at a lower field strength to be pushed toward depolarization.

## Conclusion

Non-invasive brain stimulation techniques are widely used for manipulating neuronal circuit behaviour and excitability. While the usage of these techniques is widely studied at the compartmental and whole-cell scales, less is known about the specificity of such approaches at the level of individual cells. Using models based on real pyramidal and parvalbumin neurons, we characterize their morphologies to demonstrate the observed variability in physical traits limits the specificity attainable with electric field stimulation. Our null results suggest that morphology alone may not account for the reported subtype specificity of brain stimulation paradigms and that such techniques may be limited in their specificity by morphological variability.

## Acknowledgments

We thank John Lewis and André Longtin for helpful comments.

